# Evolutionary antifragile therapy

**DOI:** 10.1101/2025.09.11.675645

**Authors:** Jeffrey West, Bina Desai, Maximilian Strobl, Jill Gallaher, Mark Robertson-Tessi, Andriy Marusyk, Alexander R. A. Anderson

## Abstract

A population of cells within a tumor can be described as antifragile (the opposite of fragile) if they derive a benefit from fluctuations or perturbations in environmental conditions induced by treatment. We hypothesized that treatment fluctuations may either promote (antifragile tumor) or inhibit (fragile tumor) the evolution of treatment resistance to targeted therapies. Analysis of the convexity of dose response curves provides a direct prediction of response to prescribed fluctuations in treatment. Theory predicts that continuous treatment protocols (i.e. zero prescribed fluctuations in treatment) will outperform uneven protocols when dose response is convex. We apply this theory to predict in vivo response to targeted therapy and validate these predictions by experimentally testing high/low intermittent dosing (uneven dosing) and continuous dosing (even dosing) schedules. Convexity (and its inverse, concavity) explains two phenomena: dose response is a convex (fragile) function but resistance onset rate is a concave (antifragile) function. Thus, we design, propose, and validate alternative treatment schedules which maximize response while maintaining prolonged sensitivity to treatment. These analyses provide supporting evidence that fluctuations alter the evolutionary trajectory of tumors in response to targeted therapy, even without altering the cumulative dose. This insight is used to design alternative, switching treatment protocols to limit the evolution of resistance, a process which we call evolutionary antifragile therapy.

## 1 Introduction

Resistance remains a paramount challenge in cancer. For example, targeted therapies show strong initial responses but rarely lead to complete tumor eradication in advanced lung cancers^1,2^, where resistance has been shown to emerge de novo from drug-tolerant persister cells leading to gradual adaptation via cooperating genetic and epigenetic adaptive changes^3^. It remains an open question whether or not we can steer or slow the adaptation of tumors to extend the efficacy of targeted treatments^4^. Below, we introduce a framework for designing treatment scheduling protocols that maximize tumor response, based on principles of antifragility^5,6^.

In times of resource scarcity or rapidly fluctuating environmental conditions, a fragile population may go extinct while robust populations survive. Robustness is considered a fundamental feature of evolvable, complex systems^7^. Functional redundancy describes robustness to an individual’s loss-of-function mutation^8^, while genotypic redundancy involves the robustness to potential environmental variation^9^.

Darwinian evolution by natural selection can be classified as an antifragile process: fragile populations tend toward extinction during adverse events, while populations that are robust (or antifragile) to fluctuations will survive. The term antifragility was originally coined to describe situations in which “some things benefit from shocks [and] thrive and grow when exposed to volatility, randomness, disorder”^5^. Darwinian evolution provides the mechanism by which populations do not just tolerate, but in fact gain a benefit from large variability in environmental conditions.

While individuals within a population may die, the population as a whole evolves toward a more antifragile state, increasingly able to cope with future perturbations on the population level^10^. The distinction between robustness and antifragility within evolutionary processes is not well-studied^11^. In cancer, it is possible to induce environmental perturbations on a prescribed protocol of timing and variation through various treatment modalities. The goal of our manuscript is to test whether tumors are fragile or antifragile to targeted therapies by direct comparison of treatment protocols with a low degree of fluctuations and a high degree of fluctuations. We hypothesize that tumors are antifragile to fluctuations in drug dosing, promoting the evolution of resistance and leading to poor outcomes for intermittent therapy. Below we review recent approaches to treatment scheduling and define terms.

### 1.1 Treatment scheduling in targeted therapy

Optimal treatment scheduling has long been studied through the integration of experimental work with theoretical, mathematical models^12–14^. Research is often focused on predictions that maximize “first-order” treatment effects, which alter the average dose delivered. For example, the “log-kill” hypothesis predicts optimality for maximum tolerated dosing with cytotoxic chemotherapy agents^15^, effectively decreasing the amount of time over which a cumulative dose is delivered, as toxicity allows^16^. Maximum tolerated dose chemotherapy schedule design often achieves only temporary response in solid tumors due to treatment resistance^17^. Alternative treatment approaches may only focus on maximizing the first-order effects of cumulative dose, such as metronomic therapy with frequent low doses known to provide an anti-angiogenic effect during chemotherapy^18,19^.

In phase 1 clinical trials for new drugs, therapy is administered continuously and uses a dose escalation protocol to find the maximal tolerated dose, which is the dose that will be used for the subsequent phases. These traditional dose selection paradigms may not apply as readily to targeted therapies that have exposure-response curves plateauing at low patient toxicities, where it is often possible to temporarily increase the dose delivered by employing intermittent off-treatment periods^4^. To name several examples, intermittent high dosing is feasible in tyrosine kinase inhibitors in HER2-driven breast cancers^20^, intermittent ribociclib (with concomitant continuous letrozole) in breast cancer^21^, intermittent erlotinib in non–small cell lung cancer^22,23^ and intermittent erlotinib once weekly in EGFR-mutant lung cancer^24^.

### 1.2 Antifragility in targeted therapy

Below, we begin by measuring an alectinib dose response curve in mice (figures 2 and 3). We show that this convex dose response curve predicts a fragile tumor response, where even treatment protocols are expected to outperform uneven protocols. However, fragility is time-dependent quantity due to a gradual onset of treatment resistance (figure 4). We show that while dose response is convex, the resistance onset is a concave function of dose. There exists a trade-off between maximizing tumor kill and maintaining treatment sensitivity, leading us to hypothesize the existence of a switching treatment schedule (even-uneven or uneven-even; figure 5) that maintains an appropriate balance within this trade-off. Mathematical modeling predicts optimal timing and experiments validate this prediction (figures 6 and 7). The theory of antifragility provides a simple and novel method to inform treatment scheduling in targeted therapy.

**Figure 1.**
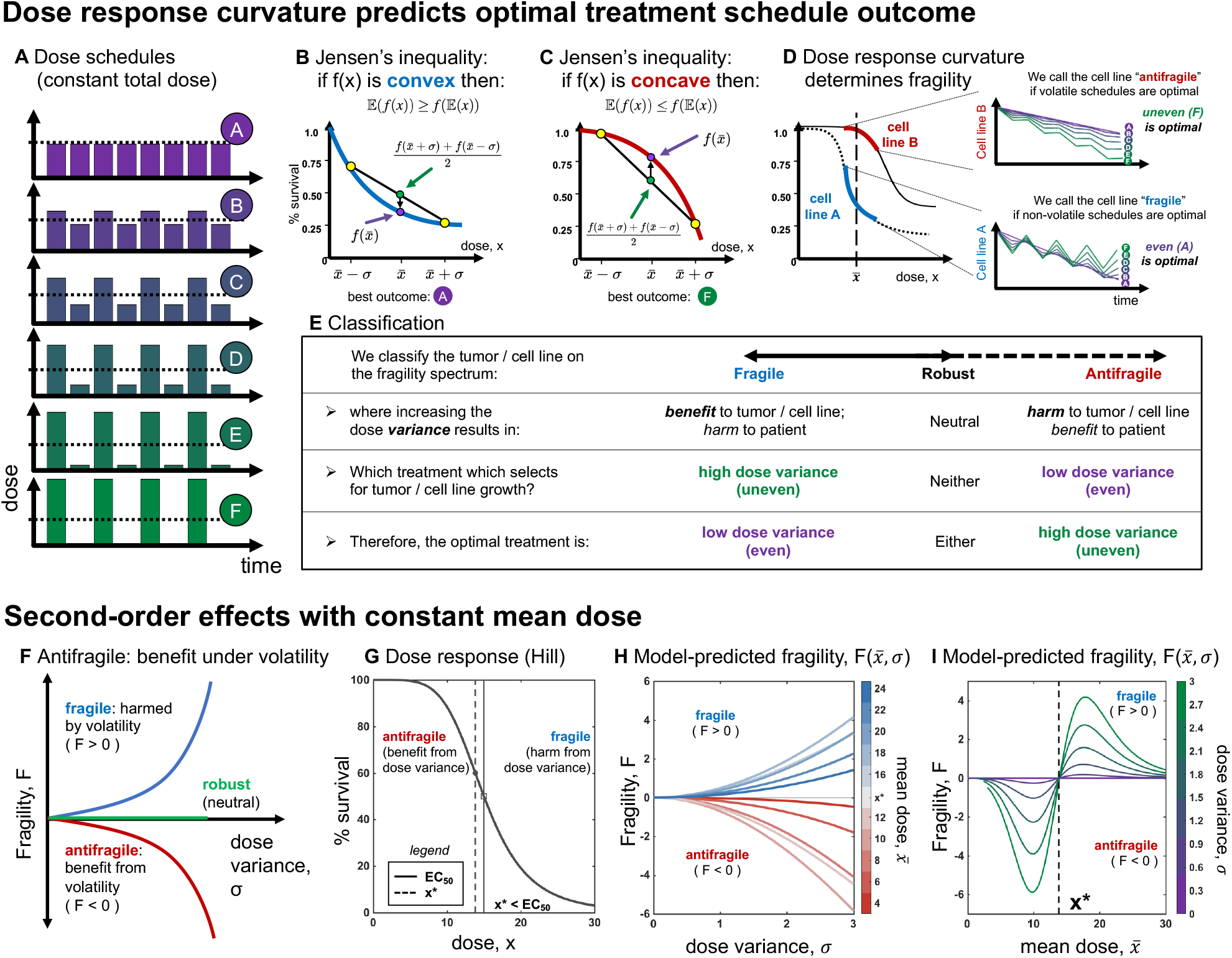
Schematic of convex and concave curvature. (A) A schematic of sample dose schedules with equivalent mean dose 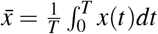. Shown for a range of dose variance, *σ*, from even (purple) to uneven (green).(B,C) Example dose response curves: convex (fragile) shown in B, and concave (antifragile) shown in C. By Jensen’s inequality, the optimal kill is achieved by even treatment if convex (B), or uneven treatment if concave (C). (D) The curvature at a given dose 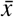, may be cell line dependent (left). Curvature predicts optimal schedule (right). (E) Classification of tumors: antifragile is a situation where dose variance causes benefit to patient, and harm to tumor. (F) Schematic of fragility, F, as a function of dose variance. (G) Example dose response (eqn. 4), delineated with antifragile (left) and fragile (right) regions. (H) Using the dose response in G, model predicted fragility as a function of variance (see eqn. 3) (I) Using the dose response in G, model predicted fragility as a function of dose mean, 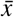 (see eqn. 3)

**Figure 2.**
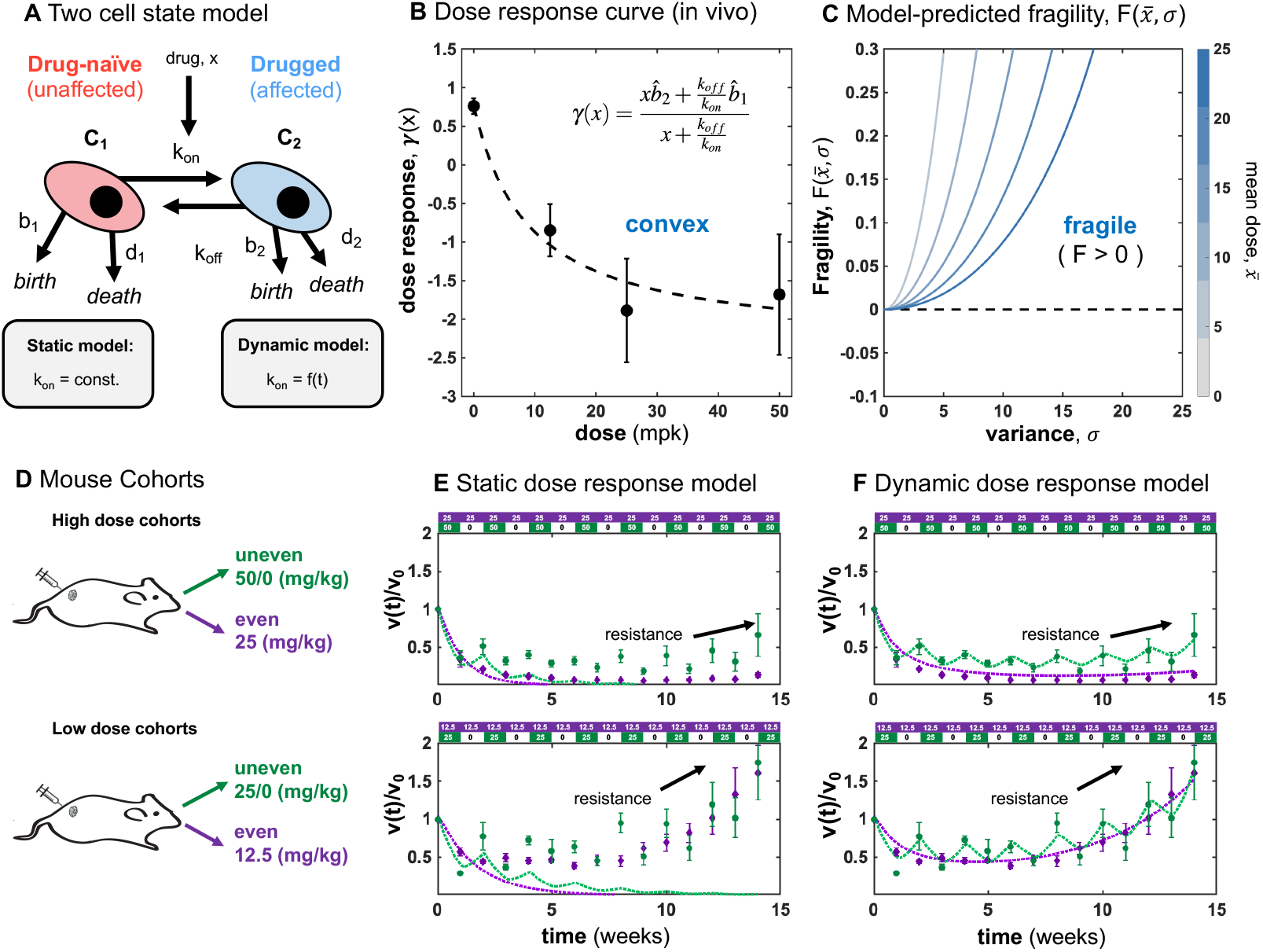
Model parameterization of high and low dose treatment with even and uneven schedules. (A) Two cell state model consisting of drug-naive cells, *C*_1_, which transition into drugged cells, *C*_2_ as a function of the drug concentration (see Methods eqn. 6 and 7). (B) In vivo dose response curve determines dosing model parameters: 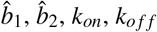 (see Methods eqn. 8). (C) Due to the convex shape of the dose response function in B, the model-predicted fragility, *F*, is fragile (*F* > 0) for the full range of dose variance (*σ* > 0) and dose mean 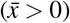. (D) Four cohorts of 6-8 mice each (see figure 3): high dose (even, uneven dosing) and low dose (even, uneven dosing) (E) Static dose response model (see Methods eqns. 6, 7 where *k*_*on*_ = const.) initially fits the data well in the first week, but fits poorly after onset of resistance. (F) Dynamic dose response model (see Methods eqns. 6, 7, with 9) improves the model fit by including *α*_*i*_, the rate of resistance onset, where the subscript *i* indicates the dose delivered each week: *i ∈* {0, 12.5, 25, 50}.

**Figure 3.**
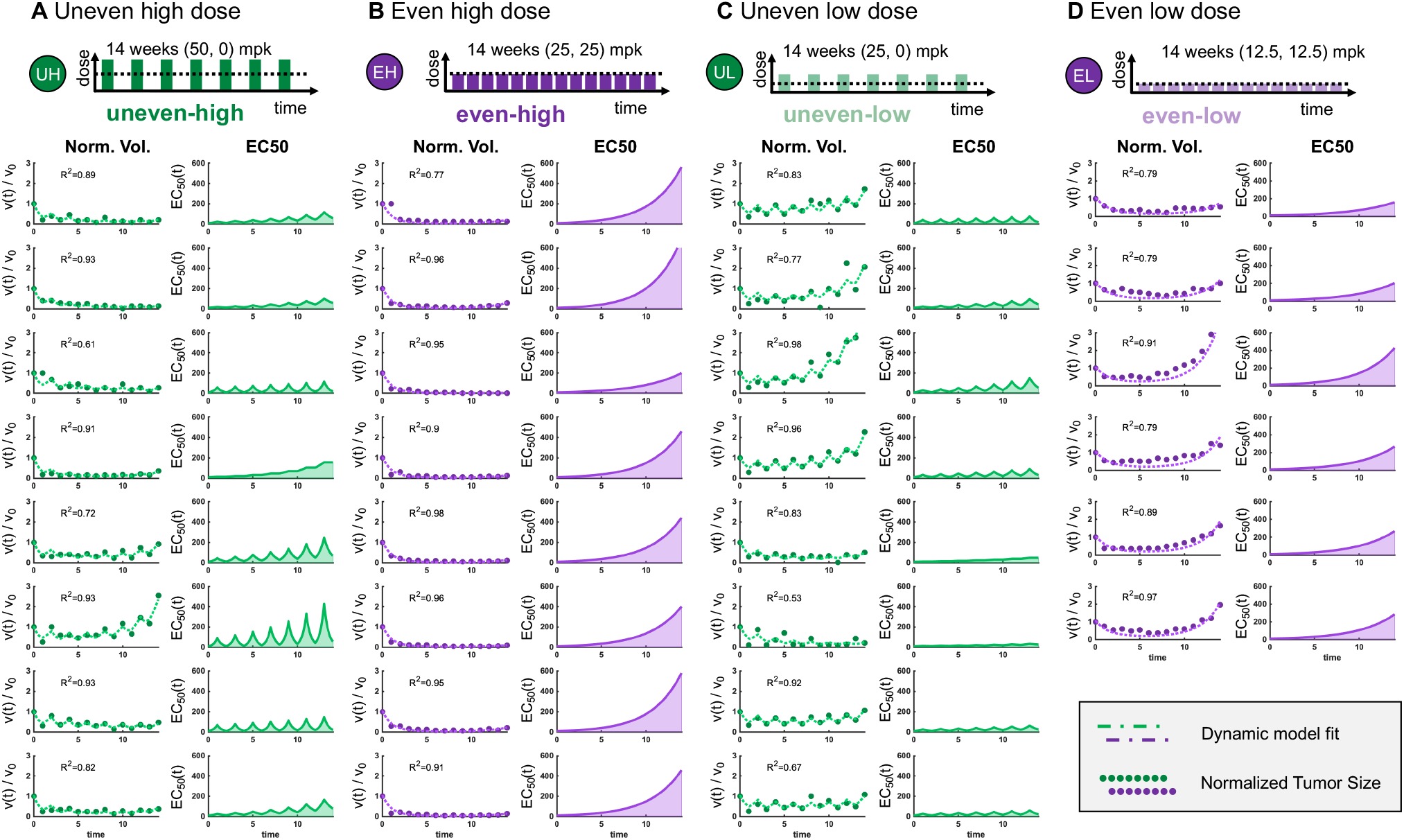
Mouse-specific parameterization of high and low dose treatment with even and uneven schedules. (A) Uneven high dose (50 mpk for one week followed by 0 mpk for one week). (B) Even, high dose (25 mpk). (C) Uneven low dose (25 mpk for one week followed by 0 mpk for one week). (D) Even, low dose (12.5 mpk). Tumor volume measurements are normalized by initial measurement, *V*_0_, and a best fit parameterization to the two cell-state model (see Methods eqns. 6, 7). Each single mouse shares dose response parameters across all cohorts 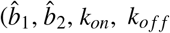; see figure 2B black dashed curve) but retains a mouse-specific value(s) for *α*_*i*_, the rate of resistance onset, where the subscript *i* indicates the dose delivered each week: *i ∈* {0, 12.5, 25, 50}.

**Figure 4.**
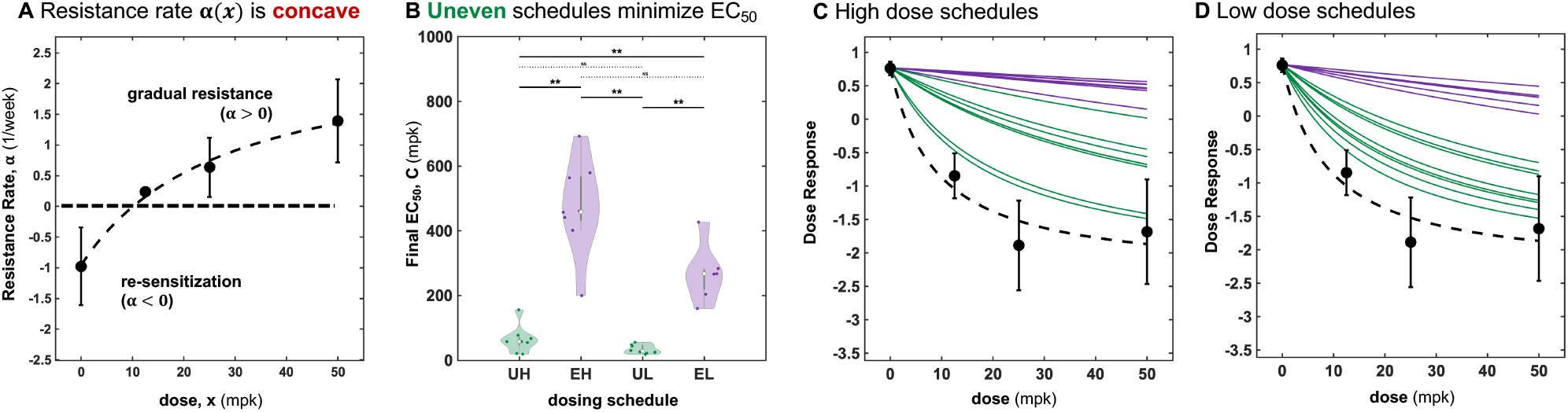
Rate of resistance is a concave function of dose. (A) Parameterization of the resistance rates *α* (see Methods eqns. 9) from figure 3 as a function of dose, *x*. The data are fit to a concave best fit (dashed line). (B) Model-predicted EC_50_ values are much lower for uneven schedules (green) than even schedules (purple) due to the concave shaped *α*(*x*). (C) Model-predicted dose response for the high dose uneven schedules (green) and even schedules (purple). (D) (C) Model-predicted dose response for the low dose uneven schedules (green) and even schedules (purple).

**Figure 5.**
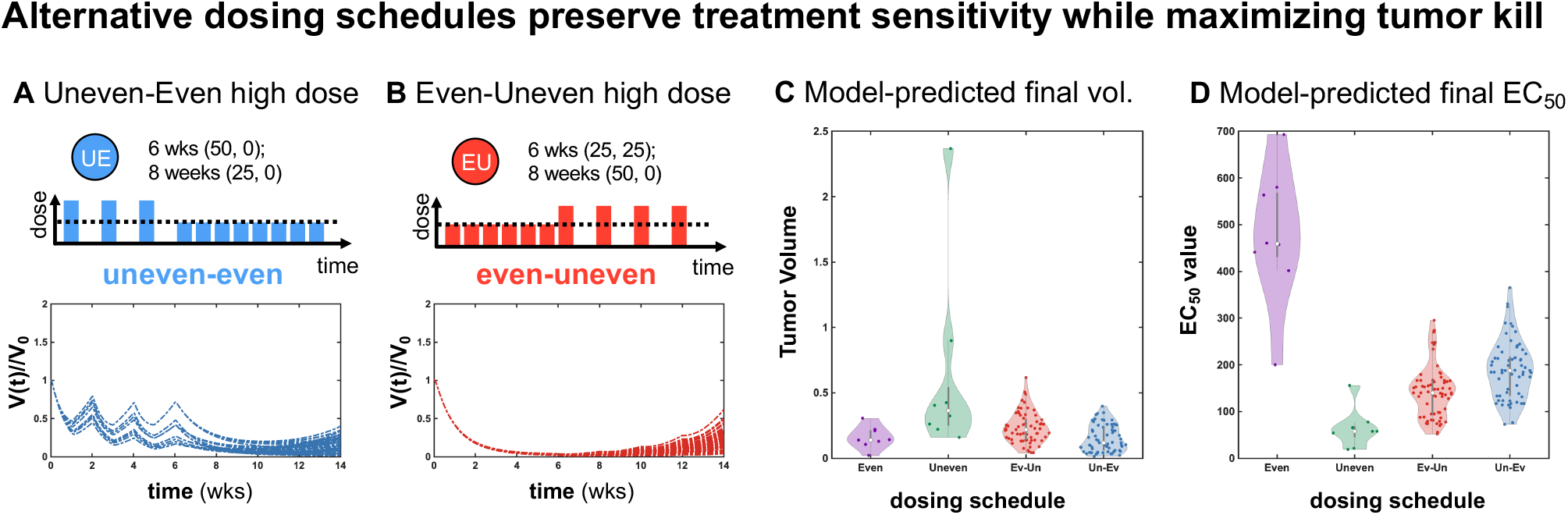
Modeling predicts that alternative dosing schedules can preserve sensitivity while maximizing tumor kill. (A,B) Based on parameterization in figure 3, two alternative schedules are simulated: Uneven dosing (6 weeks) followed by even dosing (8 weeks) shown in blue. Even dosing (6 weeks) followed by uneven dosing (8 weeks) shown in red. (C) Modeling-predicted final tumor volume. The even-uneven or uneven-even switching schedules are non-inferior to even dosing, despite maintaining a superior EC_50_ value in (D). (D) Modeling-predicted final EC_50_ value. Uneven schedules minimize EC_50_, followed by even-uneven, then uneven-even switching schedules.

**Figure 6.**
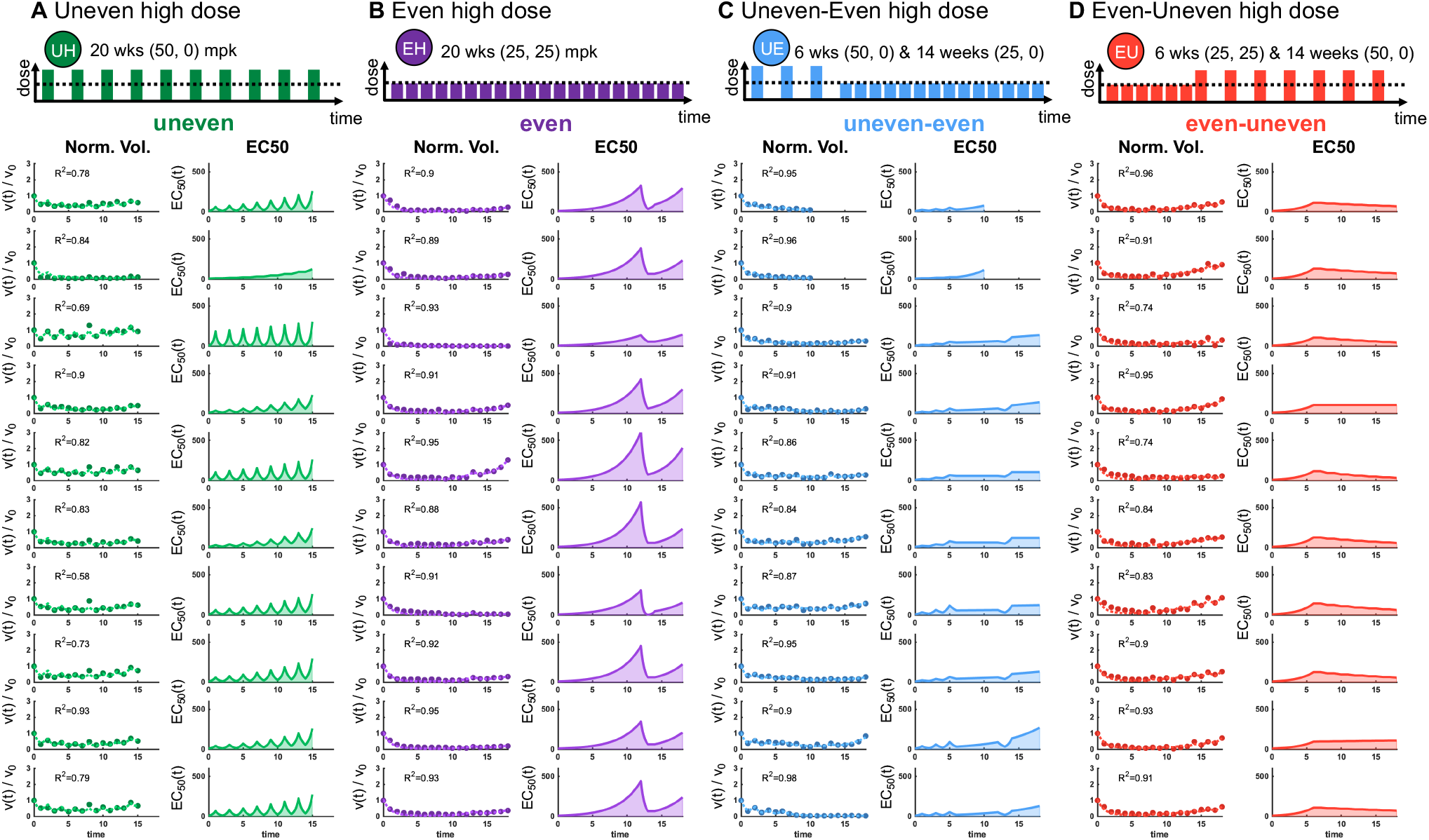
Validation experiments for proposed alternative treatment schedules. (A) Even, high dose (25 mpk) (B) Even-uneven switching: 25 mpk for six weeks followed by 50/0 mpk uneven weekly (C) Uneven-even switching: 50/0 mpk uneven for six weeks followed by 25 mpk weekly. (D) Uneven 50/0 schedule. Note: weeks 12-13 and 18-19 were administered the same {0, 50} mpk and {0, 25} mpk across all cohorts to evaluate sensitivity to same drug dose for each cohort (see figure 7).

**Figure 7.**
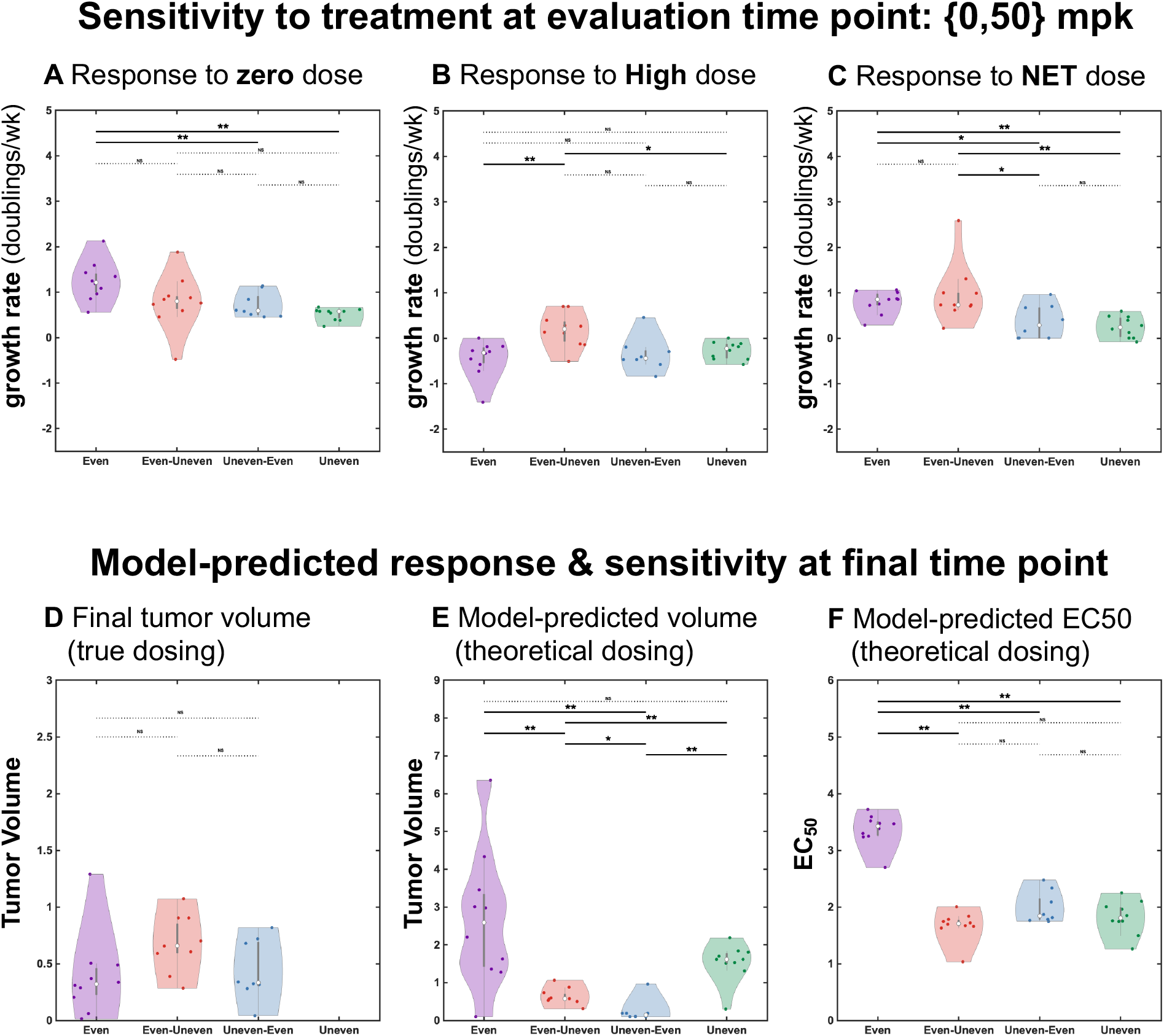
Evaluation of treatment sensitivity across treatment schedules. Top: sensitivity to treatment (proliferation rate, DIP) at evaluation time point 1 (A) Response to zero dose in week 13 (B) response to high dose (50mpk) in week 14 (C) net response to both doses across weeks 13-14. Bottom: Model-predicted response and sensitivity for theoretical dosing compared to true dosing. (D) Final normalized tumor volume, shown for the true dosing schema (including evaluation time points). (E) Model-predicted final normalized volume with evaluation time points removed and simulated for the theoretical proposed alternative schedules in 5C. (E) Corresponding model-predicted EC_50_ values for the same theoretical proposed alternative schedules.

## 2 Results

### 2.1 Dose response fragility

Improving cancer therapy will require maximizing the effectiveness of first-order effects (cumulative dose) and maximizing the effectiveness of second-order effects (the intermittent variance of dose delivered). In clinical practice, it is common to adhere to a fixed treatment protocol that administers a constant dose periodically (i.e. weekly). Alternative dosing schedules increase the dose periodically by employing intermittent off-treatment periods^21–23^. We refer to this as a “uneven” treatment schedule. We consider a treatment “cycle,” consisting of two consecutive doses, 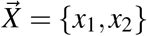, where all cycles have identical mean dose delivered such that 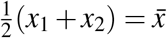. In this way, 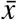 is the first-order parameter (dose mean) and *σ* is the second-order parameter (dose variance):

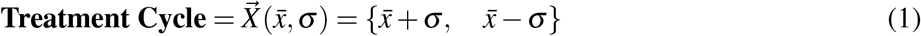

Figure 1A, depicts an even dosing schedule (purple schedule, “A”) in addition to a range of uneven schedules with increasing variance (“B” to “F”; purple to green), each with identical mean dose but differing dose variance. The outcome of these schedules can be predicted by observing the curvature of the dose response function. Examples of convex and concave dose response functions are shown in 1B and C, respectively. Let *f* (*x*) be a continuous function describing cell viability in response to dose *x*. If this function is concave (fig. 1B), the value of the dose response, 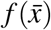, is greater than the average of the response of a high and low dose, 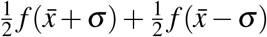. In this case, an even dosing schedule is optimal (minimizing viability). Conversely, if this function is convex (fig. 1C), the value of the dose response is less than the high/low average and the uneven dosing schedule is optimal. An explanation of this phenomena is found succinctly in Jensen’s Inequality^25,26^, which states that the expected value of a convex function is greater than the function evaluated at the expected value: 𝔼(*f* (*x*)) *≥ f* (𝔼(*x*)). The curvature predicts the optimal dose schedule for each cell line (fig. 1D right). We introduce a practical definition of fragility to compare the response of a high variance treatment cycle (*σ* > 0) with low variance (*σ* = 0):

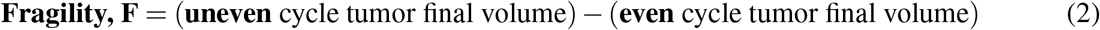

We employ the terminology “fragile” to refer to convex regions of dose response where even cycles are optimal (*F* > 0), and “antifragile” to refer to concave regions where uneven cycles are optimal (*F* < 0). In figure 1D, cell line A (blue) is fragile at dose 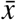, while cell line B (red) is antifragile at the same dose. Visually, fragility is determined by the curvature of the dose response: concave curvature bending downward is fragile while convex curvature bending upward is antifragile. This classification of cell lines is outlined in 1E, where antifragile cell lines should be treated with high / low dosing to maximize kill, and fragile cell lines should be treated with continuous dosing.

Shown in figure 1F, there exist three possible second-order effects: fragile (*F* > 0), robust (*F* = 0), and antifragile (*F* < 0). It is possible to precisely define fragility if the dose response function is known. For example, the bottom panels in figure 1 quantify fragility for the Hill function (fig. 1G) as a function of variance (fig. 1H) and dose (fig. 1I). The sigmoidal dose response has both convex (antifragile) and concave (fragile) regions, delineated by the inflection point of the curve.

### 2.2 Dose response inference in vivo

A parsimonious mathematical model of drug-affected and drug-unaffected cells (figure 2A) can recapitulate the *in vivo* dose response curve in figure 2B (see Methods). The y-axis measures the exponential growth rate (doublings/week) of tumor volume measured by calipers after one week of treatment. This alectinib dose response curve is characterized by a convex shape (figure 2B). A best-fit of a dose response function (Hill equation in Methods eqn. 8) is shown in the dashed lines, enabling a prediction of fragility as a function of dose mean and variance (figure 2B,C). The convex shaped dose response is fragile within the full range of dose mean and variance values. Theory predicts that even treatment protocols will outperform uneven protocols, especially at high doses.

The prediction that tumors are fragile to alectinib (and thus even treatment protocols are superior) is tested in two cohorts of mice (figure 2D), each with identical dose mean, 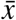. Figure 2E illustrates that the prediction based on convexity is validated, as even treatment outperforms uneven, especially at higher doses. The static mathematical model derived from the dose response curve in figure 2B (see Methods) shows a poor fit to experimental data outside of the first week (figure 2E, dashed lines), indicating the necessity to incorporate gradual resistance to treatment over time. Adding dose-dependent onset of resistance improves model fit (figure 2F, dashed lines). The dynamic dose response model incorporates a time-dependent EC_50_ value, where the convexity of the dose response may change over time.

To assess heterogeneity in dose response and resistance onset, we repeat the model fitting process for each individual mouse across all treatment conditions: 1) even treatment with a low dose 2) uneven treatment with a low dose 3) even treatment with a high dose and 4) uneven treatment with a high dose (figure 3). Uneven dosing was altered weekly (alternating with a zero dose). The structure of the mathematical model includes an initial dose response curve with corresponding EC_50_ value. The EC_50_ value increases over time at a rate dependent on the dose delivered in each treatment interval. Intriguingly, the model predicts a continuous onset of resistance for even treatment, but a period of re-sensitization (decreasing EC_50_) during off treatment periods for uneven schedules (figure 3).

### 2.3 Convexity explains response and resistance rate

Mathematical modeling provides a quantitative prediction of the resulting dose response curve after the onset of resistance. Figure 4A illustrates the rate of resistance, *α*, as a function of dose delivered. In contrast to the dose response function, the resistance rate is a concave function. Thus, theory predicts that uneven schedules minimize resistance, quantified by the model-predicted EC_50_ in figure 4B. Next, figures 4C,D plots the initial dose response curve (black dashed line) and the predicted dose response for each mouse at the final experimental time point. Uneven treatment schedules (green lines) tend to remain more sensitive to treatment than the corresponding even treatment schedules. This finding is consistent across high dose (figure 4C) and low dose (figure 4C) schedules.

Convexity (and its inverse, concavity) explains two important phenomena. First, dose response is initially convex, leading to the superiority of even treatment schedules, especially during early time points within the experimental data. Second, the emergent model-predicted rate of resistance function is concave. This concave function provides an opposite prediction: maximally suppressing the onset of resistance is achieved through uneven treatment administration.

### 2.4 Leveraging second-order effects through alternative schedules

The convex-concave response-resistance trade-off leads us to hypothesize the existence of alternative treatment schedules that may outperform fixed even or uneven dosing strategies. Two intuitive additional alternatives are proposed in figure 5C: even-uneven (red) uneven-even (blue) switching. These switching strategies aim to balance the trade-off of maximizing response (even) while maintaining good sensitivity (uneven) for prolonged time periods. Importantly, the model predicts response to treatment at the dose administered (e.g. 25mpk), but also provides a predicted response to the full range of dose values (0 through 50mpk). This allows for in silico prediction of alternative dosing schedules, such as those in figure 5C. As noted previously, we restrict ourselves to considering only schedules with identical cumulative dose, to clearly illustrate the existence of second-order effects (differential response outcomes based on dose variance only).

Mathematical modeling predicts the efficacy of these alternative strategies in figure 5A,B, using the parameterization obtained in previous figures. The model predicts similar outcomes (final volume) for even (purple) and the alternative uneven-even (blue) and even-uneven (red) dosing schedules, but predicts that both switching strategies leads to superior EC_50_ value (figure 5C,D).

### 2.5 Validation of alternative treatment strategies

Next, experimental validation of the proposed alternative schedules is shown in figure 6 for ten mice in each cohort. The proposed schedules were modified at an evaluation time point during week 12. At this time point, all four cohorts were administered the same dose of {0, 50} mpk and {0, 25} mpk, respectively, to evaluate sensitivity to an identical drug dose at an identical time point. In this manner, it is possible to validate the overall efficacy of alternative treatment schedules (final tumor volume) in addition to sensitivity (corresponding to a deeper response) to the same dose at the evaluation time points.

The same model fitting approach is undertaken: all initial mice share dose response parameters across all cohorts (see figure 5A black dashed curve) but retain a mouse-specific value for resistance onset, *α*_*i*_. Mice show some variability in response and resistance dynamics, but both the even-uneven and uneven-even schedules provide good response while maintaining a low predicted EC_50_ at the end of the experiment (figure 6B,C) when compared with their even and uneven counterparts (figure 6A,D)

### 2.6 Confounders by observations

Several conclusions can be drawn from the summary in figure 7, which delineates the proliferation rate (DIP rate) of tumors during the zero dose and high dose (50 mpk) during the evaluation time point. Even schedules are associated with a fast regrowth rate under zero dose (figure 7A), but a good response to high dose (figure 7B). In the net, the overall responsiveness during the two-week evaluation is best for the uneven schedule.

According to predictions from mathematical modeling, it is surprising that the even treatment has such a good response to a high dose in this evaluation time period. To understand this, it is important to note that the evaluation time points provide a confounding factor in accurately understanding the alternative treatment schedules. The response to a high dose is measured directly following a treatment-off period. During this short drug holiday, the model predicts a striking reduction of the EC_50_ value. Often referred to as the “observer effect,” the state of a system can be disturbed during the observation of that system. Thus, we repeated the mathematical modeling parameterization for the new round of data and removed the effect of the dose holiday during the evaluation time period. The result is shown in figure 7E, with corresponding predicted EC_50_ value in 7F, where even treatment is expected to be inferior to any other.

## 3 Discussion

Over the past decades it has become increasingly clear that the clinical benefit of a cancer therapeutic agent is determined not only by its molecular action but also by its schedule^27^. Antifragility, pioneered in financial risk management^5,26^, provides a general tool to compare treatment schedules in an intuitive yet formal fashion. Antifragility is rooted in a simple mathematical construct (the second-derivative of dose response) and is easily generalizable to a range of dose response curves obtained from experimental model systems. Dose response assays are routinely collected for *in vitro* models, but more rarely for *in vivo* or patient-derived xenograft models. Despite the widespread implementation of dose response assays in biological sciences, these curves are typically used to measure differential response in first-order effects, (mean value of drug dose delivered) while second-order effects (variance of drug dose) are generally ignored^11,26^. The benefit of variable dosing increases when the evolution of resistance to treatment occurs. We have shown that continuous administration of high doses result in similar outcomes as uneven treatment strategies, affirming non-inferiority of continuous schedules in treatment-naive settings.

This conclusion breaks down after the evolution of resistance, suggesting that treatment schedules are only optimal for limited periods of time, and will need to be adapted as the tumor changes in response to treatment. Treatment adjustments to manage toxicity are common in clinical practice, and work on so-called “adaptive therapy” has shown that personalized adjustments in dose variability are clinically feasible and beneficial^28,29^, through maintenance of drug-sensitive cells in order to suppress resistant growth due the cost to resistance^30–32^ or tumor subclonal competition^33,34^. Two main adaptive algorithms have been proposed^35^: AT1 (dose modulation) and AT2 (dose skipping). Intriguingly, recent work on PARP inhibitors in ovarian cancer has shown that the optimal adaptive algorithm depends on the curvature of the dose response function^36^. Future work can investigate if drug holidays are a viable method to push dose response toward desirable regions of antifragility. For example, adaptive administration of BRAF-MEK inhibitors in BRAF-mutant melanoma allowed for recovery of a transcriptional state associated with a low IC-50 value^37^.

It is important to note that the goal of this manuscript is to demonstrate the existence of second-order effects, and not necessarily derive a globally optimal treatment schedule. The proposed treatment strategies here are a strict subset of feasible treatment schedules, which can include altering the cumulative dose, escalation or deescalation, or other arbitrarily complicated scheduling protocols. The point of this exercise is a clear demonstration that investigation into optimizing treatment protocols requires knowledge (and measurement) of second-order effects. Here, we have shown that second-order treatment effects alter both dose responsiveness and the rate of resistance.

The framework also relies on a strong philosophical foundation. While it is difficult to calculate the patient-specific risk and timing of resistance, it is often more straightforward to predict the harm incurred if a line of treatment fails^5^.

Future work will investigate the optimal time to switch to antifragile therapy, and investigate the effects of drug synergism and antagonism. Previous theoretical work has often considered the sensitive and evolved resistant cell types in isolation, measuring the ‘ecological’ fragility based on response to dose^38,39^. Other theoretical workpredicts how these cell-cell interactions within a heterogeneous tumor influence response to second-order effects of dosing, known as ‘evolutionary fragility’^40^. Here, we have shown an in vivo, pharmacodynamic analysis of dose response curvature, but drug delivery (pharmacokinetics) will also influence the concentration over time and alter second-order effects^41^. Lastly, it is important to note that while antifragile therapy has implications on intermittent therapy, this work is not simply a reinvention of intermittent scheduling. Intermittent schedules typically utilize decreased frequency of dosing, while antifragile therapy emphasizes that dose escalation may be possible (and beneficial) when administered less frequently.

## 4 Methods

### 4.1 Xenograph studies

NCI-H3122 cells were suspended in 100*µ*l of mixture of 1:1 RPMI and BME-Type 3 (R&D Systems) and then injected 10^6^ tumor cells subcutaneously into both flanks of NOD-scid IL2Rgnull (NSG) recipient mice aged between 4 to 6 weeks. After 3 weeks post-implantation, mice were randomly allocated to receive either vehicle (water), 12.5, 25, or 50 mg/kg of alectinib, or a treatment break as per experimental needs. Tumor dimensions were assessed weekly using electronic calipers, with assumptions made about spherical tumor shape for volume calculations. The experimental procedures were conducted following the guidelines of the Institutional Animal Care and Use Committee of the H. Lee Moffitt Cancer Center. Animals were housed in a specific pathogen–free vivarium accredited by the Association for the Assessment and Accreditation of Laboratory Animal Care (AAALAC) and received care and veterinary supervision in accordance with standard guidelines, including temperature and humidity control and a 12-hour light/12-hour dark cycle.

### 4.2 Mathematical definition of (anti)-fragility

If the functional relationship between dose, *x*, and response to treatment, *f* (*x*), is known, the definition of fragility introduced in eqn. 2 can be written precisely:

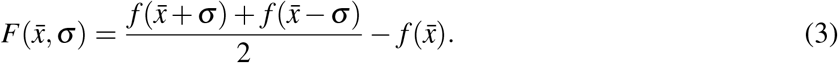

where the effect of each dose is assumed to be time-independent and response is additive. Thus, there exist three possible second-order effects: fragile (*F* > 0), robust (*F* = 0), and antifragile (*F* < 0).

Drug effect is often modeled using a Hill equation, describing the viability of a population of cells as a function of dose, *x*, can be written:

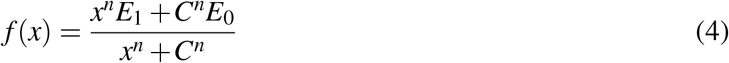

where the Hill slope, *n* represents the steepness of the Hill curve, the EC_50_ value, *C*, denotes the half-max dose concentration and *E*_0_ and *E*_1_ are minimum and maximum response values, respectively. As noted above, fragility is related to the curvature of the dose response function. Mathematically, the curvature of a function is determined by the second derivative. Thus:

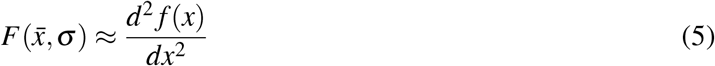

While it is straightforward to determine the second derivative of the Hill function, this only corresponds to the local curvature, this approximation is valid only for small values of dose variance, *σ* . The second derivative will produce inaccurate estimates of curvature for as dose variance increases.

### 4.3 Dose response inference

Here, fragility makes a clear prediction of treatment scheduling optimality (even versus uneven schedules) based on the dose response curve. Therefore, we need to develop a mathematical model that describes the response to dose while at the same time tracking tumor volumes over time. To do so, we use a simple two cell-state (unaffected, affected) model of drug action, as seen in reference 42. The mathematical model is a set of two ordinary differential equations describing the abundance of drug-unaffected, *C*_1_(*t*), and drug-affected, *C*_2_(*t*), cells over time.

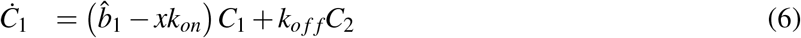

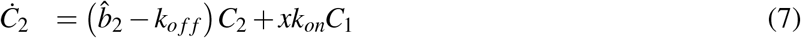

where *k*_*on*_ is the transition rate constant of the drug-unaffected to drug-affected state and *k*_*o f f*_ is the transition rate constant for the drug-affected to drug-unaffected state. Drug-unaffected and drug-affected cells grow at an exponential rate 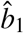 and 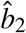, respectively. When transition rates (*k*_*on*_, *k*_*o f f*_) are large then an equilibrium between states is immediately reached, known as the partial equilibrium assumption, enabling a derivation (see ref. 42 for further details) of the drug-induced proliferation rate of the total number cells, *C*_1_ + *C*_2_.

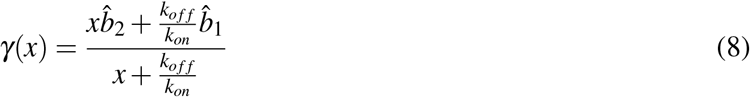

By comparison, the DIP model recapitulates a sigmoidal Hill dose response function (eqn. 4) when 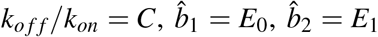, and *n* = 1. Thus, the Hill equation can be derived by considering a weighted average of drug-unaffected cells (*E*_0_), drug-affected cells (*E*_1_) operating at the equilibrium of a reversible transformation between unaffected and affected populations, which corresponding dose-dependent rates of action^43^.

### 4.4 Gradual multi-factorial resistance

The major advantage of this modeling framework is the translation between an time-varying ordinary differential equation and a static dose response curve. However, due to the onset of treatment resistance, the response to treatment is changing over time, and therefore the assumption that the effect of each dose is time-independent and response is additive is inaccurate. To account for this, we allow the unaffected-to-affected transition rate to vary with time and dose:

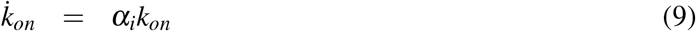

where the rate of change of *k*_*on*_ is an unknown, increasing function of dose, *α* = *α*(*x*). In practical implementation, we estimate *alpha*_*i*_ for each dose delivered, *x*_*i*_, in murine experiments, described in the next section.

### 4.5 Parameterizing experimental data

Experimental data for four cohorts of mice were used to parameterize the mathematical modeling: 1) high dose, even, 2) high dose, uneven, 3) low dose, even, and 4) low dose uneven cohorts. Each cohort consisted of six to eight mice and followed for 14 weeks administration of alectinib.

The first step in mathematical model parameterization was to take data from week 1 for all four cohorts (and untreated controls) to quantify the initial dose response curve at the beginning of treatment (see figure 2A), providing estimates for 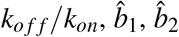 in equation 8.

The second step is to find a mouse-specific value for *α*_*i*_, the rate of resistance onset, where the subscript *i* indicates the dose delivered each week: *i ∈* {0, 12.5, 25, 50}. Thus, even treatment schedules have a single mouse-specific parameter: *α*_12.5_ (low dose even) or *α*_25_ (high dose even). In uneven treatment schedules, there are two mouse-specific parameters: *α*_25_ (low dose uneven) with corresponding off-treatment *α*_0_, or *α*_50_ (high dose uneven) with corresponding off-treatment *α*_0_. All mouse-specific parameters are averaged to give an estimate for the mean and standard deviation of resistance onset. Subsequently a best fit curve fits the mean data to infer the functional relationship between resistance and dose, *x*: *α*(*x*) (see figure 5, black dashed line).

## 5 Acknowledgments

The authors gratefully acknowledge funding by the National Cancer Institute via the Cancer Systems Biology Consortium (CSBC) U01CA232382, U54CA274507; the Physical Sciences Oncology Network (PSON) U54CA193489; and Moffitt Cancer Center support from the Center of Excellence for Evolutionary Therapy and the Cancer Biology & Evolution Program.

## 6 Competing Interests

The authors declare no competing interests.

### 7 Data Availability

The data that support the findings of this study are openly available on GitHub at:

https://github.com/MathOnco/Evolutionary-Antifragile-Therapy

## 8 Code Availability

The code used to model within this study is openly available on GitHub at:

https://github.com/MathOnco/Evolutionary-Antifragile-Therapy

## References

1. Schneider, J. L., Lin, J. J. & Shaw, A. T. Alk-positive lung cancer: a moving target. Nature cancer 4, 330–343 (2023).

2. Sasaki, T., Rodig, S. J., Chirieac, L. R. & Jänne, P. A. The biology and treatment of eml4-alk non-small cell lung cancer. European journal cancer 46, 1773–1780 (2010).

3. Vander Velde, R. et al. Resistance to targeted therapies as a multifactorial, gradual adaptation to inhibitor specific selective pressures. Nature communications 11, 2393 (2020).

4. Tracey, A. FDA’s Brian Booth: “We need to reconsider our approach to dose selection”. The Cancer Letter (2022).

5. Taleb, N. N. Antifragile: Things that gain from disorder, vol. 3 (Random House Incorporated, 2012).

6. Taleb, N. N. & Douady, R. Mathematical definition, mapping, and detection of (anti) fragility. Quantitative Finance 13, 1677–1689 (2013).

7. Kitano, H. Biological robustness. Nature Reviews Genetics 5, 826–837 (2004).

8. Láruson, Á.J., Yeaman, S. & Lotterhos, K. E. The importance of genetic redundancy in evolution. Trends Ecology & Evolution (2020).

9. Goldstein, D. B. & Holsinger, K. E. Maintenance of polygenic variation in spatially structured populations: roles for local mating and genetic redundancy. Evolution 46, 412–429 (1992).

10. Danchin, A., Binder, P. M. & Noria, S. Antifragility and tinkering in biology (and in business) flexibility provides an efficient epigenetic way to manage risk. Genes 2, 998–1016 (2011).

11. Axenie, C. et al. Antifragility in complex dynamical systems. npj Complexity 1, 12 (2024).

12. Benzekry, S. et al. Metronomic reloaded: Theoretical models bringing chemotherapy into the era of precision medicine. Seminars Cancer Biology 35, 53–61, DOI: 10.1016/j.semcancer.2015.09.002 (2015). 1011.1669v3.

13. Ledzewicz, U. & Schättler, H. Application of mathematical models to metronomic chemotherapy: What can be inferred from minimal parameterized models? Cancer Letters 401, 74–80, DOI: 10.1016/j.canlet.2017.03.021 (2017).

14. Jarrett, A. M. et al. Optimal Control Theory for Personalized Therapeutic Regimens in Oncology: Background, History, Challenges, and Opportunities. Journal Clinical Medicine 9, 1314, DOI: 10.3390/jcm9051314 (2020).

15. Skipper, H. E., Schabel, F. M. J. & Wilcox, W. S. Experimental evaluation of potential anticancer agents xiii, on the criteria and kinetics associated with” curability” of experimental leukemia. Cancer Chemotherapy Report 35, 3–111 (1964).

16. Skipper, H. E. The Effects of Chemotherapy on the Kinetics of Leukemic Cell Behavior. Cancer Research 25 (1965).

17. Group, E. B. C. T. et al. Systemic treatment of early breast cancer by hormonal, cytotoxic, or immune therapy: 133 randomised trials involving 31 000 recurrences and 24 000 deaths among 75 000 women. The Lancet 339, 1–15 (1992).

18. Miller, K. D., Sweeney, C. J. & Sledge Jr, G. W. Redefining the target: chemotherapeutics as antiangiogenics. Journal Clinical Oncology 19, 1195–1206 (2001).

19. Kerbel, R. S. & Kamen, B. A. The anti-angiogenic basis of metronomic chemotherapy. Nature Reviews Cancer 4, 423–436 (2004).

20. Amin, D. N. et al. Resiliency and vulnerability in the her2-her3 tumorigenic driver. Science translational medicine 2, 16ra7–16ra7 (2010).

21. Griffiths, J. I. et al. Serial single-cell genomics reveals convergent subclonal evolution of resistance as patients with early-stage breast cancer progress on endocrine plus cdk4/6 therapy. Nature Cancer 1–14 (2021).

22. Chmielecki, J. et al. Optimization of dosing for egfr-mutant non–small cell lung cancer with evolutionary cancer modeling. Science translational medicine 3, 90ra59–90ra59 (2011).

23. Schöttle, J. et al. Intermittent high-dose treatment with erlotinib enhances therapeutic efficacy in egfr-mutant lung cancer. Oncotarget 6, 38458 (2015).

24. Grommes, C. et al. “pulsatile” high-dose weekly erlotinib for cns metastases from egfr mutant non-small cell lung cancer. Neuro-oncology 13, 1364–1369 (2011).

25. Jensen, J. L. W. V. et al. Sur les fonctions convexes et les inégalités entre les valeurs moyennes. Acta mathematica 30, 175–193 (1906).

26. Taleb, N. N. (anti) fragility and convex responses in medicine. In International Conference on Complex Systems, 299–325 (Springer, 2018).

27. Strobl, M., Gallaher, J., Robertson-Tessi, M., West, J. & Anderson, A. Treatment of evolving cancers will require dynamic decision support. Annals Oncology 34, 867–884 (2023).

28. Zhang, J., Cunningham, J. J., Brown, J. S. & Gatenby, R. A. Integrating evolutionary dynamics into treatment of metastatic castrate-resistant prostate cancer. Nature Communications 8, 1816 (2017).

29. Gatenby, R. A., Silva, A. S., Gillies, R. J. & Frieden, B. R. Adaptive therapy. Cancer Research 69, 4894–4903 (2009).

30. West, J. et al. Towards multi-drug adaptive therapy. Cancer Research (2020).

31. Gallagher, K. et al. Mathematical model-driven deep learning enables personalized adaptive therapy. Cancer Research (2024).

32. West, J. et al. A survey of open questions in adaptive therapy: Bridging mathematics and clinical translation. Elife 12, e84263 (2023).

33. Strobl, M. A. et al. Turnover modulates the need for a cost of resistance in adaptive therapyturnover modulates resistance costs in adaptive therapy. Cancer Research 81, 1135–1147 (2021).

34. Viossat, Y. & Noble, R. J. The logic of containing tumors. bioRxiv (2020).

35. Enriquez-Navas, P. M. et al. Exploiting evolutionary principles to prolong tumor control in preclinical models of breast cancer. Science Translational Medicine 8, 327ra24–327ra24 (2016).

36. Strobl, M. A. et al. To modulate or to skip: De-escalating parp inhibitor maintenance therapy in ovarian cancer using adaptive therapy. Cell Systems (2024).

37. Smalley, I. et al. Leveraging transcriptional dynamics to improve braf inhibitor responses in melanoma. EBioMedicine 48, 178–190 (2019).

38. West, J. et al. Antifragile therapy. BioRxiv 2020–10 (2020).

39. Taleb, N. N. & West, J. Working with convex responses: Antifragility from finance to oncology. Entropy 25, 343 (2023).

40. Bayer, P. & West, J. Games and the treatment convexity of cancer. Dynamic Games Applications 13, 1088–1105 (2023).

41. Pierik, L., McDonald, P., Anderson, A. R. & West, J. Second-order effects of chemotherapy pharmacodynamics and pharmacokinetics on tumor regression and cachexia. Bulletin Mathematical Biology 86, 47 (2024).

42. Harris, L. A. et al. An unbiased metric of antiproliferative drug effect in vitro. Nature Methods 13, 497–500 (2016).

43. Meyer, C. T. et al. Quantifying drug combination synergy along potency and efficacy axes. Cell Systems 8, 97–108 (2019).

